# Tirzepatide preserves hematopoietic stem and progenitor cycling while remodeling inflammatory monocytes in obese mice

**DOI:** 10.64898/2026.09.01.748392

**Authors:** Nathan M. Krah, Elena Gonzalez-Alvarado, Amog P. Urs, Chinmayee Goda, Yaphet Bustos, James Marvin, Bradley D. Weaver, Spencer Gygi, Ashish Toshniwal, Dennis Towne, Alvaro Jesus Narbona-Perez, Katarina Heyden, Juan A. Cantres-Velez, Corey N. Cunningham, Sankalp Arora, Ramiro Garzon, Jared Rutter, Adrienne M. Dorrance, Amandine Chaix

## Abstract

Obesity expands myeloid progenitors, myelopoiesis and increases the production of monocytes. While weight loss (WL) alleviates aspects of this inflammatory dysregulation, it is not known whether GLP-1 receptor agonists or other traditional modalities of WL differentially modify hematopoietic stem/progenitor cells (HSPCs), hematopoiesis, or inflammatory cell production. To test this, we compared the hematopoietic compartment in lean, obese and weight-reduced mice from tirzepatide treatment and caloric restriction (CR) implemented to match the body weight in both groups. At equal WL, we found CR induced multilineage cytopenias, whereas tirzepatide preserved blood lineages while specifically reducing classical Ly6C^hi^ CCR2^+^ monocytes. To define the mechanisms underlying these changes we performed single-cell mRNA sequencing of bone marrow HSPCs and mature mononuclear blood cells. CR-HSPCs suppressed gene sets associated with nutrient sensing, proliferation and oxidative phosphorylation (OXPHOS) and exhibited lower inferred cell cycle activity, whereas tirzepatide-HSPCs attenuated these changes. Unlike CR, we found that across progressively differentiated cells from HSPCs to mature blood monocytes, tirzepatide increasingly suppressed OXPHOS and simultaneously shifted the maturation spectrum away from classical monocytes. Following six weeks of tirzepatide withdrawal and weight regain, Ly6C^hi^ CCR2^+^ monocytes rebounded to levels seen in obese mice. These findings suggest that tirzepatide uncouples WL from the broad hematopoietic suppression seen in CR by preserving progenitor activity but selectively remodeling inflammatory/classical monocytes. We demonstrate that WL modality differentially impacts hematopoietic adaptation and provide evidence that classical monocytes are an effector cell through which tirzepatide may dampen obesity-associated inflammation.

**Key Points:**

- At equivalent weight loss, calorie restriction causes cytopenias and suppresses HSPC cycling, while tirzepatide preserves these parameters
- Tirzepatide reduces inflammatory monocytes, shifts maturation, decreases OXPHOS genes, and monocytes rebound after drug withdrawal

## Introduction

Obesity induces myeloid-biased hematopoiesis and high fat diet (HFD) activates hematopoietic stem and progenitor cells (HSPCs), expanding myeloid progenitors, leading to increased production of monocytes and macrophages^1,2^. Inflammation from peripheral tissues feeds back on bone marrow (BM) progenitors to amplify and sustain myelopoiesis and monocytosis^3,4^. This feed-forward loop of myelopoiesis generates a sustained source of inflammatory cells that persists even after weight loss (WL) and contributes to progressive chronic inflammation and cardiometabolic disease^5^. Thus, the hematopoietic system represents an important upstream driver of the chronic inflammation and associated disease risk in obesity. Understanding whether hematopoietic “memory” of obesity persists after WL and whether different forms of WL alter this source of inflammatory cells is of high clinical importance. Yet relatively little is known about how different forms of WL remodel BM HSPCs and their progeny.

The topic of WL modality has become increasingly important over the last five years, as new treatment options have emerged for patients with obesity. For example, the dual GLP-1/GIP receptor agonist, tirzepatide, induces substantial WL and improves cardiometabolic outcomes in obesity^6,7^. Additionally, emerging evidence suggests that GLP-1R agonists exert broad anti-inflammatory effects that occur independently of WL ^8–11^. Several studies suggest that these medications reduce circulating inflammatory mediators and modulate monocyte and macrophage inflammatory states^12,13^. But whether GLP-1-based therapies act upstream to remodel HSPC activity, differentiation and hematopoietic output remains largely unknown. Given recent increases in GLP-1 use, defining whether these medications alter obesity-associated inflammatory hematopoiesis, and if they do so differently than conventional WL modalities, is a timely and clinically relevant question.

To address this, we integrated peripheral blood phenotyping with single-cell mRNA sequencing (scRNA-seq) of BM HSPCs and mature blood cells from weight-reduced mice to assess the effect of WL on hematopoiesis. Importantly, we compared tirzepatide-treated to weight-matched calorie restricted (CR) mice to determine whether the WL modality differentially impacted hematopoietic output. We used this approach to determine whether equivalent reductions in body weight produce divergent hematopoietic responses and identify the distinct effects of tirzepatide and CR on HSPC gene programs and mature myeloid-cell states.

## Materials and Methods

### Mice and dietary interventions

Male C57BL/6J mice (JAX 000664), 8 weeks of age, were fed a 60% kcal high-fat diet (Research Diet D12492) *ad libitum* for 12 weeks to induce obesity. Obese mice were then randomized to: (i) continued *ad libitum* HFD; (ii) tirzepatide (3 nmol/kg BW, daily subcutaneous injections) with *ad libitum* HFD; or (iii) weight-matched calorie restriction, achieved by daily restriction of HFD to match the body weight of the tirzepatide group. A separate age-matched lean control cohort was maintained on *ad libitum* low-fat diet (6 % kcal fat; Inotiv 2920X). Body weight was recorded at least once per week. For the tirzepatide and CR withdrawal experiment, interventions were discontinued at maximal stable WL, ∼10 weeks after treatment initiation, and mice were followed for an additional 6 weeks with access to *ad libitum* HFD and we performed longitudinal bleeds as described below. All animal experiments were carried out in accordance with the guidelines and approved by the University of Utah Institutional Animal Care and Use Committee (IACUC).

### Complete blood counts and immunophenotyping/flow cytometry

Peripheral blood was collected in EDTA pre-coated tubes. Complete blood counts were measured on Heska Element HT5 (Antech Diagnostics, Inc). Immunophenotyping was performed by the University of Utah flow cytometry core facility. Briefly, red blood cells were lysed with ammonium chloride solution for 5 minutes and samples were spun at 400 x g for 5 minutes. Cells were then resuspended in 200 μl of PBS with BSA. Surface antibody cocktail was added (**Table S1**) and incubated for 45 minutes at room temperature. Cells were washed, fixed and permeabilized with eBioscience Foxp3/Transcription factor buffer set according to the manufacturer’s recommendations. Following permeabilization, Foxp3 antibody was added and incubated for 45min and then washed. Data was acquired on a Cytek Aurora spectral cytometer (Cytek Biosciences). The gating scheme to define relevant populations was designed as follows: cleanup gates consisted of forward scatter area and side scatter area along with forward scatter width to define single cells. Following this, cells were gated on CD45 and Zombie UV to characterize live immune cells. Next, major subsets of cells were defined by CD3 (T-cells), NK1.1 (NK cells), B220 (B-cells), Ly6G (neutrophils), and Siglec F (Eosinophils). T-cells were further separated into gamma delta T-cells, FoxP3 Tregs, CD4 conventional T-cells, and CD8 T-cells. The remaining myeloid cells were separated into PDCA-1^+^ DCs as well as monocytes, which were CD11b^+^ CX3CR1^+^. Finally, classical and non-classical monocytes were identified based on Ly6C and CCR2 expression.

### Single-cell RNA sequencing

Peripheral blood mononuclear cell (PBMC) libraries were each prepared from two pooled mice (n=3 libraries, i.e., 6 mice, per group), whereas CD117-enriched marrow libraries were prepared from individual mice (n=3 libraries/mice per group). Accordingly, the biological replicate unit is the library level, and all blood/PBMC statistics were computed at the library level. CD117 (c-Kit)-enriched bone marrow cells were isolated from the right femur and tibia, as well as the left femur of all mice, and anatomic sampling was consistent across all dissections. Anti-CD117 microbeads (Miltenyi Biotec, cat. no. 130-091-224) were used to enrich progenitors. CD117-enriched marrow cells and PBMCs were profiled with the 10x Genomics GEM-X Flex v2 gene expression assay in 4-plex format (GEM-X Flex Mouse Transcriptome Probe Kit v2, PN-1000902; GEM-X Flex Sample Preparation v2 Kit, PN-1000781; GEM-X Flex Gene Expression Chip Kit, PN-1000791; Dual Index Kit TS Set A, PN-1000251; GEM-X Flex v2 Mouse, PN 1000931; **Table S2**), following user guide CG000834. We prepared libraries for 12 PBMC and 12 CD117-enriched marrow samples (3 libraries per treatment group) according to the manufacturer’s protocol and minimized batch effects by multiplexing samples using a balanced block design; each 4-plex library was demultiplexed by probe barcode into four per-sample matrices. Sequencing was performed by the University of Utah Genomics Core Facility on an Illumina NovaSeq X (paired-end 151 x 151 cycles; NovaSeq X Series 25B reagent kit, PN 20104706) to a target depth of 50,000 reads per cell.

### Read processing, quality control, integration, and annotation

Reads were aligned and quantified with Cell Ranger multi version 10.0.0 using the 10x Chromium Mouse Transcriptome Probe Set v2.0.0 and the GRCm39 reference (10x build 2024-A). For our analysis, we retained all cells that had >200 detected genes and <10% mitochondrial reads (**Table S3**). We did not apply an upper unique molecular identifier (UMI) bound and did not perform computational doublet removal. Counts were normalized to 10,000 per cell and log1p-transformed. Using Seurat flavor, we selected 2,000 highly variable genes per library, scaled to a maximum of 10, and took 50 principal components. We integrated the libraries using Harmony. A 15-nearest-neighbor graph was built on the Harmony embedding, cells were clustered with Leiden (igraph, undirected, 2 iterations, resolution 1.0), and UMAPs were generated. We assigned clusters to cell types using previously defined canonical markers with Wilcoxon rank-sum marker detection (scanpy rank_genes_groups) used to surface cluster-enriched genes and guide assignments (**Table S4a-d**)^14,15^.

To analyze peripheral blood monocytes, these cells were re-embedded separately (1,500 highly variable genes, Harmony, Leiden resolution 0.6). Sub-clusters were scored with marker modules to stratify cells as either classical (Ly6C^hi^/Ccr2^+^), intermediate, or non-classical (Ly6C^lo^/Ccr2^-^) monocytes. We treated maturation as continuous and used diffusion pseudotime (scanpy diffmap/dpt) on the Harmony-integrated monocyte graph. The marker used in our flow cytometry studies, *Ly6c2*, could not be used here because it does not have a probe in the Flex set. To circumvent this, the root was set to the cell with the highest mean expression of the three well-characterized classical monocyte markers *Ccr2*, *Vcan* and *Sell*, and the axis was oriented so pseudotime increased toward the non-classical state, which was checked against *Nr4a1* expression.

### Pseudobulk differential expression and gene-set enrichment

UMI counts were summed within each population to one pseudobulk profile per library and modeled in edgeR with quasi-likelihood fits^16^. We then tested the following contrasts for differential gene expression: CR vs. HFD, tirzepatide vs. HFD, tirzepatide vs. CR, and HFD vs. LFD. As expected, the number of differentially expressed genes varied by contrast but there were no individual genes that passed FDR adjustment for tirzepatide vs. HFD (**Tables S5, S6**). Given limited differential gene expression, we focused on transcriptomic inferences at the gene-set and model level.

Pre-ranked enrichment was run with fgsea v1.28.0^17^, based on the GSEA framework^18^, against MSigDB Hallmark^19^, C2:CP:Reactome and C5:GO:BP collections, retrieved for the appropriate species (*Mus musculus*) through msigdbr v26.1.0. Each contrast was ranked using two different methods: first by signed significance, sign(log2FC) * −log10(*p*), with *p* floored at the smallest non-zero value, and second, by log2 fold-change alone. We ran fgsea under both rankings (minSize 10, maxSize 500, eps 0). A pathway was called concordant and significant if it was enriched in the same direction and passed a Benjamini-Hochberg adjusted *p* < 0.05 under both rankings (**Tables S7, S8a–b**).

To confirm the HSPC gene set results in Figure 2, we ran three robustness analyses (**Figure S3**). First, we re-ranked genes by the signed edgeR quasi-likelihood statistic, sign(log2FC) x √F, and repeated fgsea. Second, we applied CAMERA, an established competitive gene-set test that accounts for intergene correlation^20^ and repeated the analysis. Finally, we restricted our analysis to a “core” HSC subset defined by a curated HSC-stemness signature^21^. The top third of cells with the highest expression of this signature was used to define core HSCs; we rebuilt pseudobulk, and re-tested. CR-associated suppression persisted with all three analyses.

### Module and cell cycle scores

Normalized log1p-transformed counts were used to compute module scores with scanpy score_genes. We scored five gene sets, including: Hallmark oxidative phosphorylation, Hallmark MYC targets v1, Hallmark E2F targets, Reactome respiratory electron transport, and GO:BP oxidative phosphorylation (**Table S9**). We assigned cell-cycle phase using scanpy score_genes_cell_cycle with the appropriate mouse orthologs of the gene sets defining S and G2/M phase by Tirosh and colleagues.^22^ The cycling fraction was scored as the proportion of cells that were not in G1.

We modeled per-library module scores for the OXPHOS, MYC and E2F gene sets in HSC/MPP, cMoP and blood classical monocytes in R as score ∼ group x compartment. The treatment x compartment interaction was tested as a difference-in-differences contrasting the effects of CR and tirzepatide between HSC/MPP and blood classical monocytes, using an ordinary linear model with libraries treated as independent replicates (n=3 per treatment per compartment). This interaction was robust to leave-one-library-out refitting. We also fit the three-compartment omnibus model with library as a random intercept, to account for HSC/MPP and cMoP scores which were measured from the same marrow library (p = 0.022; Table S10). We repeated the analysis with the Reactome and GO:BP OXPHOS gene sets, as well as the Hallmark MYC and E2F targets, to confirm our findings.

To ask whether the tirzepatide-mediated OXPHOS effect was independent of the cell maturation shift, per-cell scores were fit as OXPHOS ∼ maturation + group + (1|library) in lmerTest, with library as a random intercept. We then evaluated group contrasts at matched maturation using Satterthwaite degrees of freedom.

### Receptor transcript detection

To determine if *Glp1r* and *Gipr* were detectable in our dataset, we scored transcript detection as the percentage of cells in a defined population with at least one raw UMI for a transcript. We reasoned that low detection could be due to either true low expression or a technical artifact. To help address this, we tested a previously published mouse pancreatic β-cells scRNA-seq dataset (Cell Ontology “type β pancreatic cell”, 140,984 cells, 10x 3′ v2/v3) retrieved from the CZ CELLxGENE Discover Census through cellxgene-census^23^. The β-cell dataset used 3′ chemistry rather than Flex and was utilized only as a detectability control and not for any quantitative comparisons.

### Statistics

Tests and corrections for each panel are described in the figure legends. All reported p-values are two-sided, with p < 0.05 used to determine significance. Unless otherwise stated in the figure legend, all plotted data are presented as mean ± SEM.

One-way ANOVA with Tukey’s multiple comparisons test was performed for all figures relating to complete blood counts and flow cytometry in Figure 1. For the tirzepatide and CR treatment withdrawal experiment described in Figure 5, within-group difference from baseline was tested using paired t-tests. To assess specificity of change between groups, we tested the group x time interaction. This interaction was tested by one-way ANOVA on per-animal change scores (Δ = post − pre) followed by Tukey HSD post hoc comparisons.

**Figure 1:**
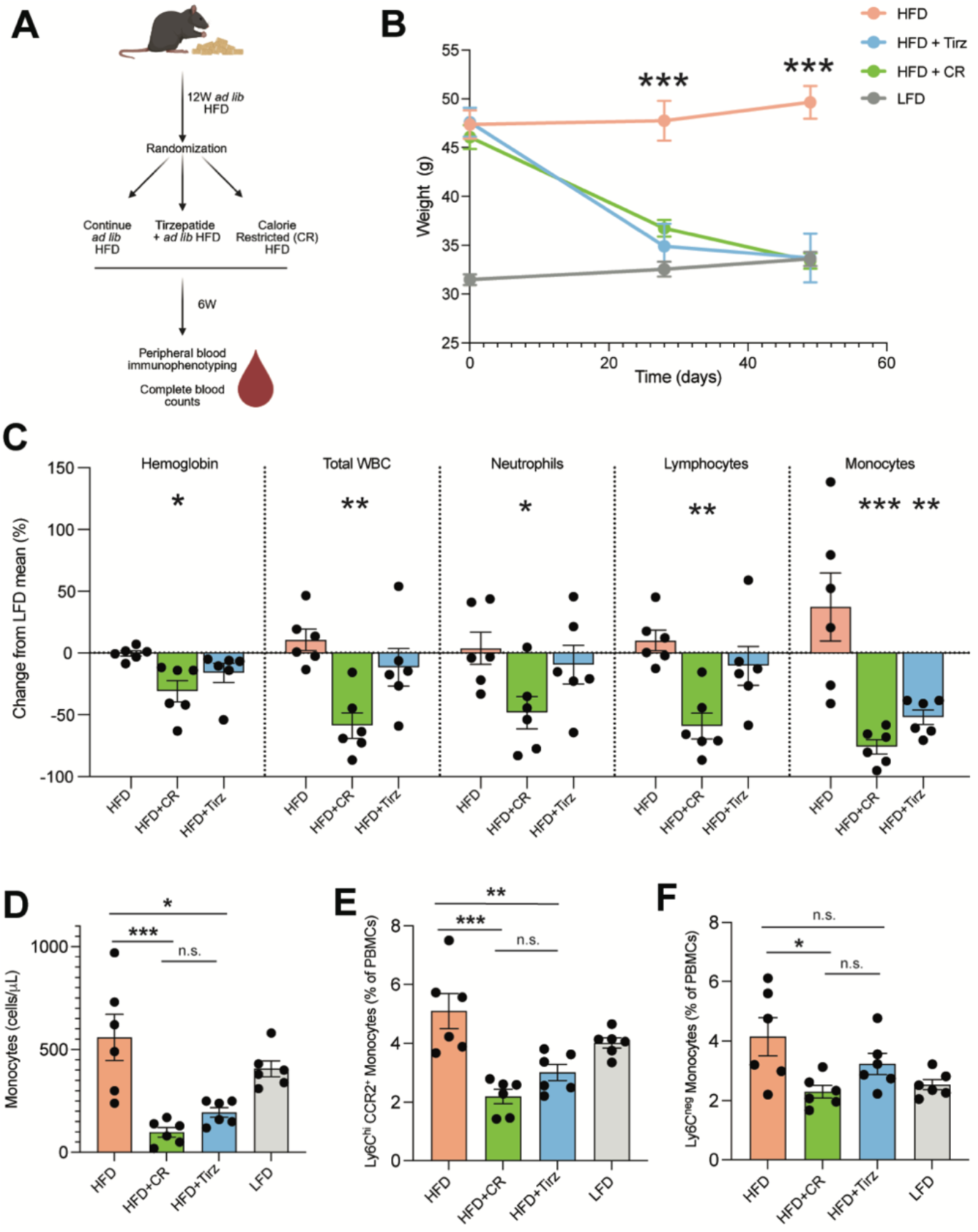
At equal WL, CR and tirzepatide produce differing hematologic phenotypes in HFD-induced obesity. **(A)** Study design: mice were fed an *ad libitum* high-fat diet (HFD) for 12 weeks to induce obesity before being randomized to continue *ad libitum* HFD, receive tirzepatide with access *ad libitum* HFD, or weight-matched CR with HFD. Mice fed *ad libitum* LFD served as lean comparators for the duration of the study. Peripheral blood complete blood counts (CBCs) and immunophenotyping were performed 6 weeks after treatment initiation (n=6 per group). **(B)** Body weight over time. Symbols represent the group mean ± SEM. **(C)** CBC parameters (hemoglobin, total white blood cell count, neutrophils, lymphocytes, and monocytes) expressed as percent change from the LFD group mean. Statistical comparisons are among HFD, HFD+CR, and HFD+Tirz, and asterisks denote comparison to HFD. **(D)** Absolute blood monocyte count (cells/µL). **(E)** Classical Ly6C^hi^ CCR2^+^ monocytes and **(F)** non-classical Ly6C^neg^ monocytes as a percentage of total PBMCs. Bars represent the mean ± SEM; symbols are individual mice, n=6 per group. Statistics were performed as one-way ANOVA with Tukey’s multiple comparisons test. \**p*<0.05, \*\**p*<0.01, \*\*\**p*<0.001; n.s., not significant.

**Figure 2:**
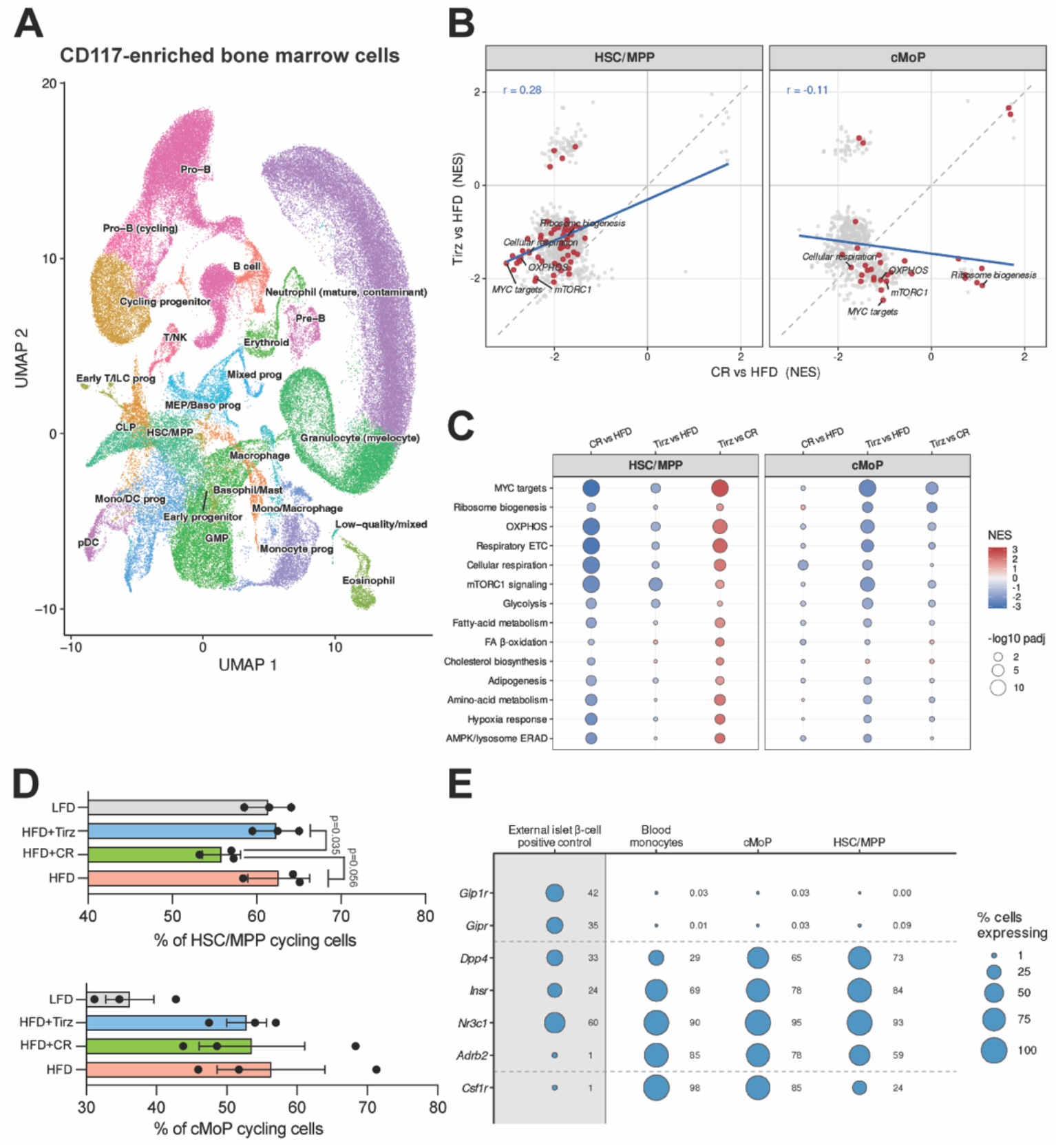
The divergence of CR and tirzepatide hematopoietic phenotypes originates in progenitors and is differentiation stage specific. **(A)** Single-cell RNA-seq of CD117-enriched HSPCs (n=3 mice/libraries per group) was performed and resulting UMAP of the integrated CD117-enriched marrow dataset with annotated populations is shown. **(B)** Genome-wide pathway response: each point represents a gene set plotted by normalized enrichment score (NES) for CR vs. HFD (x-axis) and tirzepatide vs. HFD (y-axis), in either HSC/MPP (left) and cMoP (right). Pathways related to nutrient-sensing and cell proliferation are highlighted in red. Pearson correlation is shown. **(C)** GSEA dot plot of curated nutrient-sensing and proliferation pathways across three indicated contrasts in HSC/MPP and cMoP compartments; dot color represents scaled NES and the dot size is scaled to the −log10 adjusted *p*-value. **(D)** Estimated fraction of cycling HSC/MPPs or cMoPs in the S/G2M phase; bars represent mean ± SEM. Symbols are individual mice/libraries (*p*=0.035 for CR vs. Tirz and *p*=0.056 for CR vs HFD by one-way ANOVA followed by Tukey’s multiple-comparisons test). **(E)** Dot plot indicated percentage of cells expressing the indicated GLP-1 or myeloid-associated transcripts. An external β-cell dataset served as a transcript-detectability control for *Glp1r* and *Gipr* (gray shaded column). Dot size and printed value signify the percentage of cells with detectable transcript.

**Figure 3:**
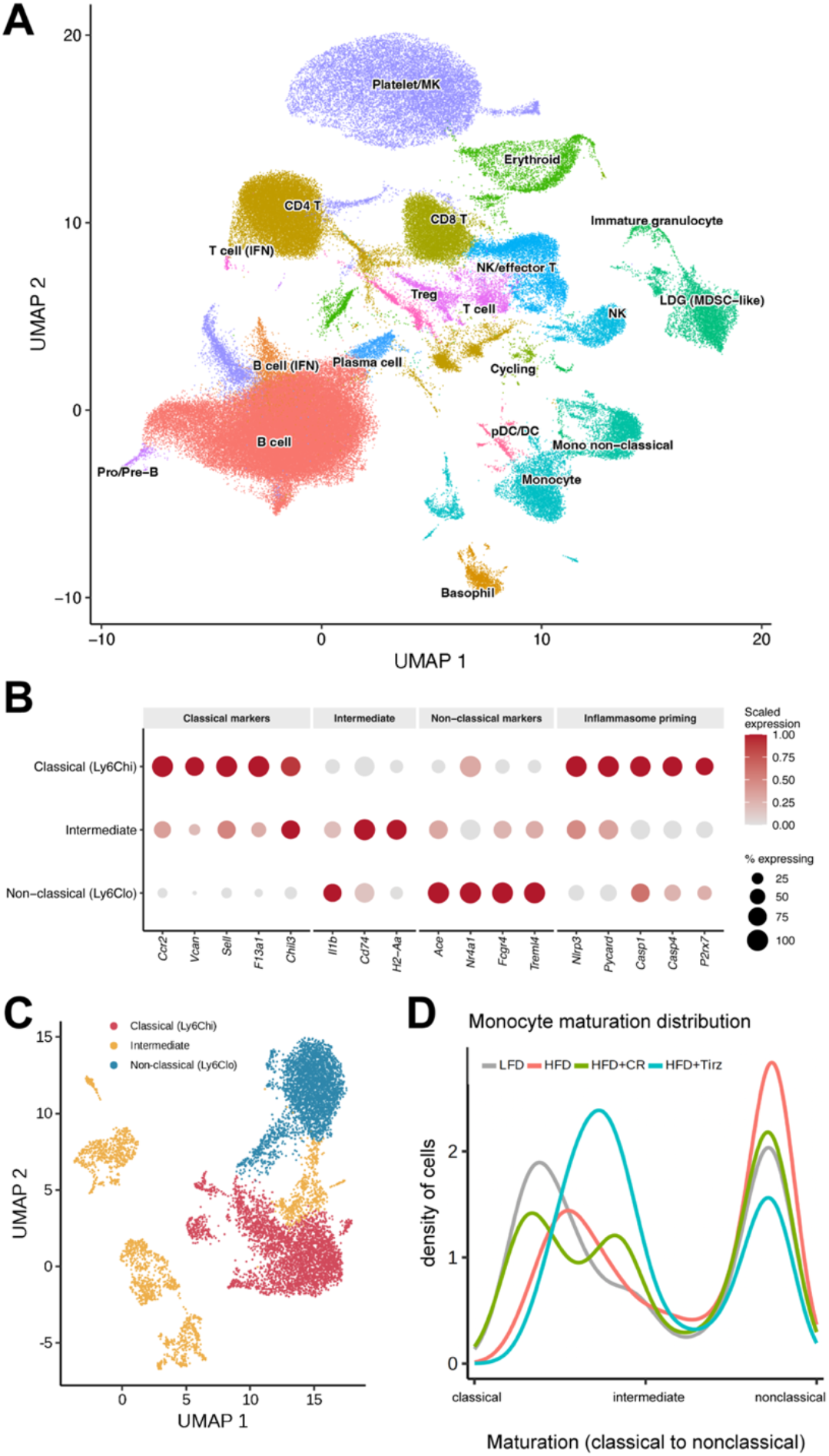
Single-cell mRNA sequencing of PBMCs identifies a full continuum of monocytes and demonstrates that tirzepatide shifts monocytes toward an intermediate phenotype. **(A)** Complete UMAP of the integrated PBMC dataset with annotated cell types, showing the complete spectrum of blood cells. **(B)** Dot plot of canonical genes defining monocyte subtypes, including inflammasome-associated genes, which are found exclusively in the classical subtype. Dot color represents expression for each gene, while dot size represents the percentage of cells expressing the indicated transcript. **(C)** UMAP of peripheral blood monocytes showing the spectrum of classical (Ly6C^hi^), intermediate and non-classical (Ly6C^lo^) differentiating monocytic phenotypes. **(D)** Diffusion pseudotime-defined distribution of blood monocytes along the classical-to-non-classical maturation trajectory. Density curves are shown by treatment group. A left or right shift is indicative of an altered distribution. PBMC libraries were generated from pooled blood of two mice (n=3 libraries per group, n=6 mice per group).

**Figure 4:**
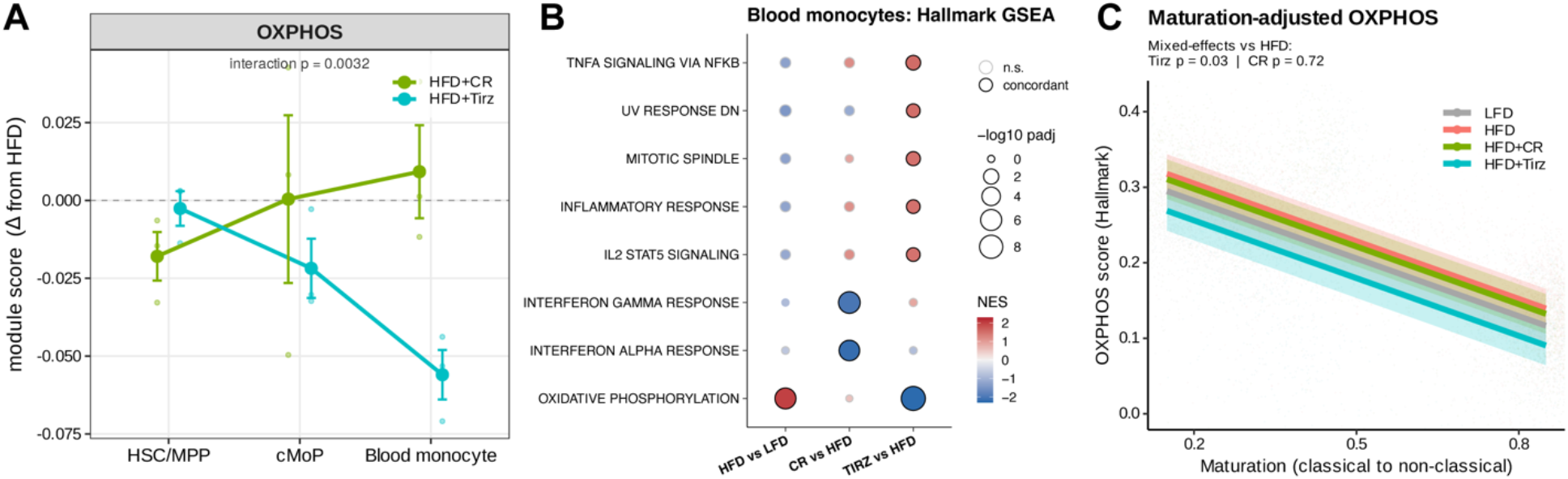
Tirzepatide-associated OXPHOS suppression emerges with monocytic maturation. **(A)** Treatment by compartment interaction: the change in OXPHOS module score compared to HFD across the maturing compartments including HSC/MPP, cMoP, and classical monocytes for HFD+CR (green) and HFD+Tirz (blue). The reported interaction is from the prespecified 2 x 2 CR-versus-tirzepatide x HSC/MPP-versus-blood classical-monocyte model (*p* = 0.0032); cMoP is shown to illustrate the differentiation trajectory. Smaller symbols are individual libraries with the large symbols representing the group mean ± SEM. **(B)** Hallmark GSEA of peripheral blood monocytes across three indicated contrasts. Dot color represents the normalized enrichment score (NES), and dot size represents the -log10 adjusted *p*-value. The bold outlines represent concordance under both ranking metrics, while a faint outline is not significant. **(C)** OXPHOS module score across monocyte maturation states in all four treatment groups, indicated by color. Lines are fitted values from a linear mixed-effects model, and the shaded areas indicate 95% confidence intervals.

**Figure 5:**
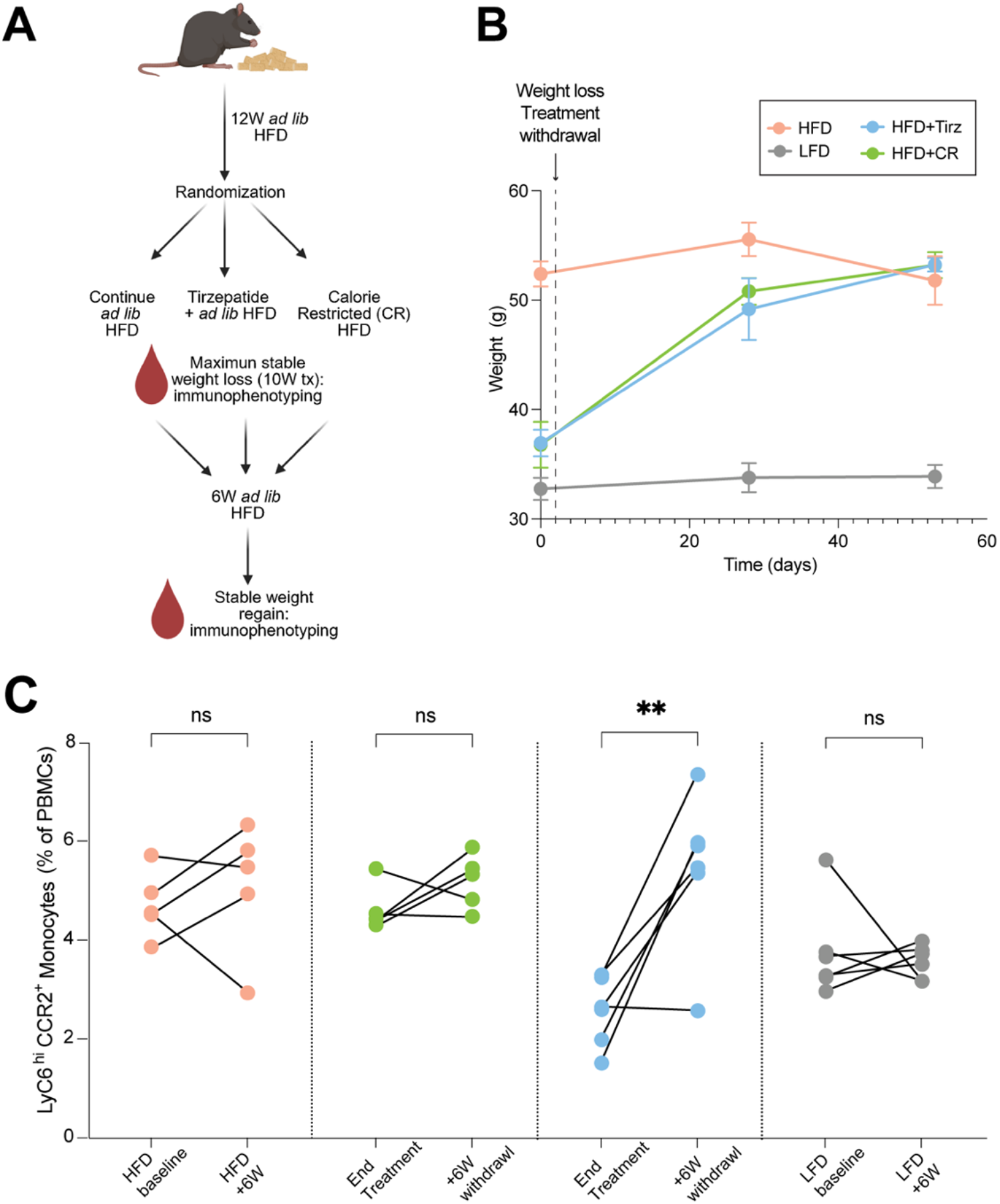
Classical monocytes rebound after tirzepatide withdrawal and weight regain. **(A)** Experimental schematic depicting overall study design: mice were fed HFD for 12 weeks to induce obesity before being randomized to continue *ad libitum* HFD alone, *ad libitum* HFD while receiving tirzepatide, or CR on a HFD to match the weight of the tirzepatide arm for 10 weeks. At stable WL body weight, samples were drawn for peripheral blood immunophenotyping. Treatment was then discontinued and blood was drawn 6W later for the same studies. Continuously fed LFD mice served as lean controls for the duration of the study. **(B)** Body weight after CR or tirzepatide withdrawal (n=6 per group). **(C)** Longitudinal Ly6C^hi^ CCR2^+^ monocytes as a percent of total PBMCs, paired within each mouse, at the end of treatment vs. 6 weeks with access to *ad libitum* HFD (n=5-6 per group). Measurements are paired in individual mice. The group x time interaction was significant (*p* = 0.004), and the post-weight gain increase in the tirzepatide group exceeded the changes observed in every other group (Tukey-adjusted *p*=0.004 versus LFD, *p*=0.025 versus HFD, and *p*=0.041 versus CR). \*\**p*<0.01; n.s., not significant.

For scRNA-seq experiments, single-cell measurements were averaged to library-level prior to statistical testing (n=3 per group). Where possible we avoided treating individual cells as independent biological replicates. As stated above, GSEA determined significance using Benjamini-Hochberg FDR. We report GSEA enrichment as adjusted q-values. All OXPHOS module scores were compared to HFD using uncorrected Welch t-tests. Electron transport chain complexes were compressed into a single composite per sample by first z-scoring each complex across all samples, then averaging across complexes within sample (Figure S4D). This composite score was compared using a one-way ANOVA. Complexes are displayed individually in Figure S4C, however they were not powered sufficiently to find significance.

### Software

Python v3.11.15 (scanpy v1.11.5, anndata v0.12.19, numpy v2.4.4, pandas v2.3.3, scipy v1.17.1, statsmodels v0.14.6), R v4.3.3 (edgeR v4.0.16, fgsea v1.28.0, msigdbr v26.1.0, lmerTest v3.1.3, emmeans v1.10.0, car v3.1.2, data.table v1.14.10, ggplot2 v3.4.4), and GraphPad Prism v11.

## Results

C57BL/6J male mice were fed a 60% HFD for 12 weeks before being randomized to continue *ad libitum* HFD without intervention, tirzepatide treatment with access to *ad libitum* HFD, or weight-matched CR with HFD (**Figure 1A**). Mice on *ad libitum* low-fat diet (LFD) served as a lean reference group. Tirzepatide and CR produced nearly identical WL by 4 weeks and matched the LFD group by the end of week 7 (**Figure 1B**). Despite nearly equivalent body weight, mice on tirzepatide and CR had different hematologic outputs. CR caused broad cytopenias, reducing hemoglobin by 31%, total white blood cells by 63%, neutrophils by 50%, and lymphocytes by 63% relative to HFD while these lineages were not affected in the tirzepatide group (n.s. vs. HFD for each) (**Figure 1C**). CR and tirzepatide both reduced peripheral monocytes (84% and 62% respectively, **Figure 1C and 1D**), but in a subtype-specific manner. While classical Ly6C^hi^ CCR2^+^ monocytes were reduced in both groups (**Figure 1E**), only CR suppressed nonclassical Ly6C^neg^ monocytes (**Figure 1F**). Taken together, at equivalent WL, CR and tirzepatide produce distinct hematopoietic phenotypes with CR inducing broad cytopenias that are absent in tirzepatide-treated mice.

We next wanted to determine how tirzepatide treatment preserves hematopoiesis during WL. To do this we performed scRNA-seq on CD117-enriched bone marrow HSPCs and peripheral blood mononuclear cells (PBMCs) from mice in all four treatment groups. After QC, cell recovery, genes detected per cell, UMI depth, and mitochondrial read fractions were comparable between treatment groups. Integrated embedding showed intermixing of cells from individual libraries and treatment groups in both datasets (**Figure S1**, **Tables S3, S4**).

Our CD117-enriched dataset resolved the expected cell lineages including HSC/multipotent progenitors (MPPs), lineage-restricted progenitors, including common monocyte progenitors (cMoP), and many maturing populations (**Figure 2A, S2**). At the pathway level, we found striking divergences in transcriptional responses between CR and tirzepatide treatment (**Figure 2B-C**). In HSPCs, CR suppressed proliferative and biosynthetic programs including MYC targets, E2F targets, G2M checkpoint, oxidative phosphorylation, and mTORC1 associated programs (**Figure 2B-C, Tables S6, S8)**. Changes in these programs were weaker in HSPCs from tirzepatide-treated animals (**Figure 2B-C**). These findings withstood edgeR quasi-likelihood statistical reranking, CAMERA gene-set testing, and restriction to an HSC-enriched subset of cells (**Figure S3A-B**). In contrast to HSPCs, tirzepatide suppressed MYC and metabolic-related pathways to a greater extent in cMoPs, while CR had a minimal effect on these pathways in this compartment (**Figure 2B-C**). To assess whether these pathway-level changes reflected reduced HSPC proliferation, we estimated the fraction of HSPCs assigned to S/G2M and found cell cycling was lowest in CR animals (55.8%) compared with HFD (62.6%), tirzepatide (62.3%), and LFD (61.4%) (**Figure 2D**). Continuous S-phase and G2/M scores showed the same pattern (**Figure S3C**). This difference was not reproduced in cMoPs, where cycling fractions were similar between HFD, CR and tirzepatide-treated animals (**Figure 2D**). Taken together, these data suggest that CR broadly suppresses HSPC activity, providing insight into the cytopenias observed with this form of WL (**Figure 1**).

To begin determining whether the effects seen at the pathway level were direct, we measured the drug target transcripts, *Glp1r* and *Gipr*, which were nearly undetectable in all lineages examined (**Figure 2E**). To exclude lack of probe coverage as an explanation for their near-zero detection, we validated that both *Glp1r* and *Gipr* were represented in the 10x Flex panel. To further support our ability to detect these transcripts if present, we analyzed an external mouse β-cell dataset in which *Glp1r* and *Gipr* were robustly detected in ∼40% of cells.^23^ While we cannot completely rule out low-level expression, our data suggest that tirzepatide acts indirectly on myeloid cells *in vivo*, aligning with recent studies suggesting that incretins temper inflammation via indirect mechanisms^9–11^.

Given that tirzepatide induced transcriptional changes in cMoPs but did not decrease their predicted cycling activity, we reasoned changes in monocytes could become more pronounced as these cells further differentiated. To test this, we performed scRNA-seq on PBMCs and this integrated dataset resolved major blood cell populations, including monocytes and dendritic cells (**Figure 3A**). After isolation of monocytes, these cells subclustered into classical, intermediate and non-classical states (**Figure 3B-C**). Monocyte states were further supported by expression of canonical genes, and only the classical/inflammatory subsets expressed high levels of individual inflammasome components (**Figure 3B, Table S9**). Because classical Ly6C^hi^ monocytes give rise to intermediate and nonclassical cells, we asked whether WL alters this maturation distribution^24^. Compared with other groups, tirzepatide shifted the monocyte maturation spectrum away from the classical phenotype and toward the intermediate state (**Figure 3D**), consistent with the reduction in classical Ly6C^hi^ CCR2^+^ monocytes measured by flow cytometry (**Figure 1E**).

Analysis across the differentiation trajectory from HSPC, cMoP and mature monocytic compartments revealed opposing trajectories specifically of the OXPHOS transcriptional program. Suppression of OXPHOS with CR was greatest in the HSC/MPPs, while tirzepatide suppressed OXPHOS progressively in more differentiated cells (**Figure 4A, Figure S4A, Table S11**). This compartment-specific interaction was replicated in three different OXPHOS-associated gene-set definitions (**Figure S4B**). To further validate these findings, unbiased Hallmark GSEA of total blood monocytes identified OXPHOS as the dominant program suppressed with tirzepatide treatment (**Figure 4B, Table S7**). We found similar reductions in gene programs related to inner mitochondrial membrane electron transport complexes I-IV and ATP synthase in monocytes from tirzepatide-treated animals (**Figure S4C-D**). Together, these data support that tirzepatide induces a coordinated metabolic remodeling in monocytes, rather than acting on a single pathway definition.

Because OXPHOS activity varied with monocyte maturation, we queried whether the tirzepatide-associated low OXPHOS state could be an artifact of this cell composition change. In a mixed effects model, monocytes from tirzepatide-treated animals retained significantly lower OXPHOS scores than HFD monocytes, whereas CR did not differ from HFD (**Figure 4C**). These data suggest that tirzepatide-mediated suppression of the OXPHOS gene program occurs throughout the maturation spectrum and is not simply explained by changes in monocyte phenotype composition.

Recent studies suggest that cessation of tirzepatide causes rebound in multiple cardiometabolic parameters related to weight regain^25^. To begin testing whether this applied to the hematologic parameters measured in our study, we asked whether reductions of classical/inflammatory monocytes persisted after discontinuation of WL interventions. After 12 weeks of HFD-induced obesity, a separate cohort of HFD, LFD, HFD+CR and HFD+Tirz mice were treated to maximal WL for 10 weeks, before discontinuation of WL interventions. We performed longitudinal peripheral blood immunophenotyping after 10W of intervention, and again 6 weeks after treatment discontinuation (**Figure 5A**). Following cessation, mice previously treated with tirzepatide or CR regained substantial body weight (**Figure 5B**). We longitudinally profiled Ly6C^hi^ CCR2^+^ monocytes and found they increased from 2.6% at maximal WL, to 5.5% of PBMCs 6 weeks after tirzepatide withdrawal (+113%), whereas there was no significant within-group change in HFD, CR, or LFD (**Figure 5C**). The magnitude of this change differed across groups (group x time interaction *p*=0.004), and the increase in monocytes after tirzepatide withdrawal significantly exceeded changes in every other group (Tukey-adjusted *p*=0.004 vs. LFD, *p*=0.025 vs HFD, and *p*=0.041 vs CR). This selective rebound of inflammatory monocytes following tirzepatide discontinuation is suggestive of a treatment-dependent phenotype that is lost upon phenotypic reversion.

## Discussion

Together, our findings identify divergent hematopoietic responses to equivalent WL with CR and tirzepatide treatment. Consistent with previous studies, our results support a model where CR restrains proliferative and metabolic programs in HSPCs limiting proliferation and inducing quiescence^26,27^. By contrast, tirzepatide preserves HSPC cycling, but exerts its strongest effect in mature inflammatory monocytes, where it decreases gene programs associated with OXPHOS and the number of circulating cells. Beyond hematopoiesis, the tirzepatide-associated suppression of OXPHOS could have consequences on monocyte function. Previous studies have shown that mitochondrial electron transport supports NLRP3 inflammasome activation^28^. Additionally, a recent study demonstrated that OXPHOS and mitochondrial electron transport are required for monocyte chemotaxis and monocyte-to-macrophage cell differentiation^29^. Although we did not directly measure these parameters, our findings are consistent with this paradigm and suggest that oxidative metabolism is a potential link between tirzepatide treatment and inflammatory monocyte behavior.

Perhaps most importantly, our weight-matched study design uncouples the effects of incretin therapy and CR-induced WL on hematopoiesis and suggests that hematologic remodeling is not determined only by the magnitude of WL. Our findings suggest that the modality of WL determines the hematopoietic response and is consistent with studies demonstrating that obesity-induced changes in HSPCs and myeloid output are not normalized by WL alone^5^. We extend this concept by showing that two interventions producing equivalent WL can generate distinct hematopoietic and inflammatory states. Going forward, it will be interesting to test whether either WL modality reverses the epigenetic changes seen in HSPCs in obesity^30^.

Following tirzepatide treatment withdrawal, classical monocytes rebounded toward levels observed in obese mice, paralleling the rebound of other cardiometabolic clinical parameters seen in humans after treatment discontinuation^25^. This argues against a durable reprogramming of cMoPs and other progenitors and suggests that suppression of monocytes depends on the systemic state of tirzepatide treatment. The factors responsible remain unknown but could include changes in nutrient availability, insulin sensitivity, or adipose-tissue inflammation among other candidates.

Our results have implications beyond obesity-associated monocytosis. In humans, incretins reduce cardiovascular events and improve heart failure outcomes in obese individuals, and WL is unlikely to fully explain these benefits^31,32^. In animal models, incretins reduce atherosclerosis and decrease the number of macrophages at disease sites, independent of WL^33^. Clonal hematopoiesis of indeterminate potential (CHIP) is independently associated with atherosclerotic cardiovascular disease and mechanistically, inflammasome function is required for CHIP-mediated atherosclerosis^34,35^. Furthermore, obesity promotes clonal expansion in CHIP^36^. Our findings therefore introduce the hypothesis that incretin-based therapies could modify the detrimental effects of myeloid cells/monocytes in CHIP patients. Direct testing will be required to determine if incretins influence clonal fitness, inflammatory monocyte number/behavior, or the cardiovascular consequences of CHIP.

Our study has limitations. All analyses were performed in male mice, and whether sex modifies the hematopoietic response to CR or tirzepatide remains unknown. This is particularly relevant to the clinic where incretin use for WL in both clinical trials and real-world patients are majority female^6,37,38^. Some of the phenotypes we identified, such as tirzepatide-dependent decreases in monocytic OXPHOS, reflect transcriptional pathway activity rather than direct measurements. Finally, our inability to detect *Glp1r* or *Gipr* suggests that there are indirect effects on myeloid cells, but we cannot rule out low-level expression. Tirzepatide-mediated systemic signals are likely to modify monocytes and identifying those signals and how they modify the function of myeloid cells will be important priorities for future studies.

Overall, our study demonstrates that CR and tirzepatide-induced WL produce fundamentally different hematopoietic and inflammatory states. While CR causes broad cytopenias and decreases HSPC cycling, tirzepatide tempers this phenotype while reversibly suppressing classical monocytes. These data suggest that WL modality influences hematopoietic adaptation and indicate that tirzepatide modifies obesity-associated inflammation by modifying classical monocytes.

## Acknowledgements

We thank additional members of the A.C. laboratory, including undergraduate research assistants Cristina Miyasato, Seth Roberts, and Jackson Henderson and PhD students Renan F.L.Vieira, Peyton D Mower and Sawyer R Sanchez for assisting with the weight loss study daily feeding and injections. We acknowledge the University of Utah Flow Cytometry Core for use of equipment and thank the staff for their assistance. Research reported in this publication also used the High Throughput Genomics Core Facility at Huntsman Cancer Institute (Illumina NovaSeq X; managed by B. Dalley) and the Cancer Bioinformatics (RNA sequencing alignment and analysis; C. Stubben). Shared resources at Huntsman Cancer Institute at the University of Utah are supported by NCI P30CA042014. NMK is funded by the NIH/NHLBI, grant *F32HL177926* and a generous Young Investigator Award from the Joe W. and Dorothy Dorsett Brown Foundation. AC thanks the University of Utah College of Health and the Center for Metabolic Health for their support in this area of research. This work was partly supported by National Institutes of Health grants UH2CA286584 and P30DK020579 to A.C., and University of Utah Vice President of Research FY25 Seed to EGA.

## Author contributions

Conceptualization: NMK and AC

Investigation: NMK, EGA, APU, CG, YB, BDW, SG, AT, DT, AJNP, KH, JACV, CNC, SA

Formal Analysis: NMK, EGA, APU, CG, BDW, DT

Data Curation: NMK, EGA, APU, CG, BDW, DT

Visualization: NMK

Resources: NMK, RG, JR, AMD, AC

Writing, Original Draft: NMK and AC

Writing, Review & Editing: All authors

Supervision: NMK, AMD, AC

Project Administration: NMK, AC

Funding Acquisition: NMK, RG, JR, AMD, AC

