## Supplementary Figures 1-4 for "Tirzepatide preserves hematopoietic stem and progenitor cycling while remodeling inflammatory monocytes in obese mice"

### Supplementary Figure 1:

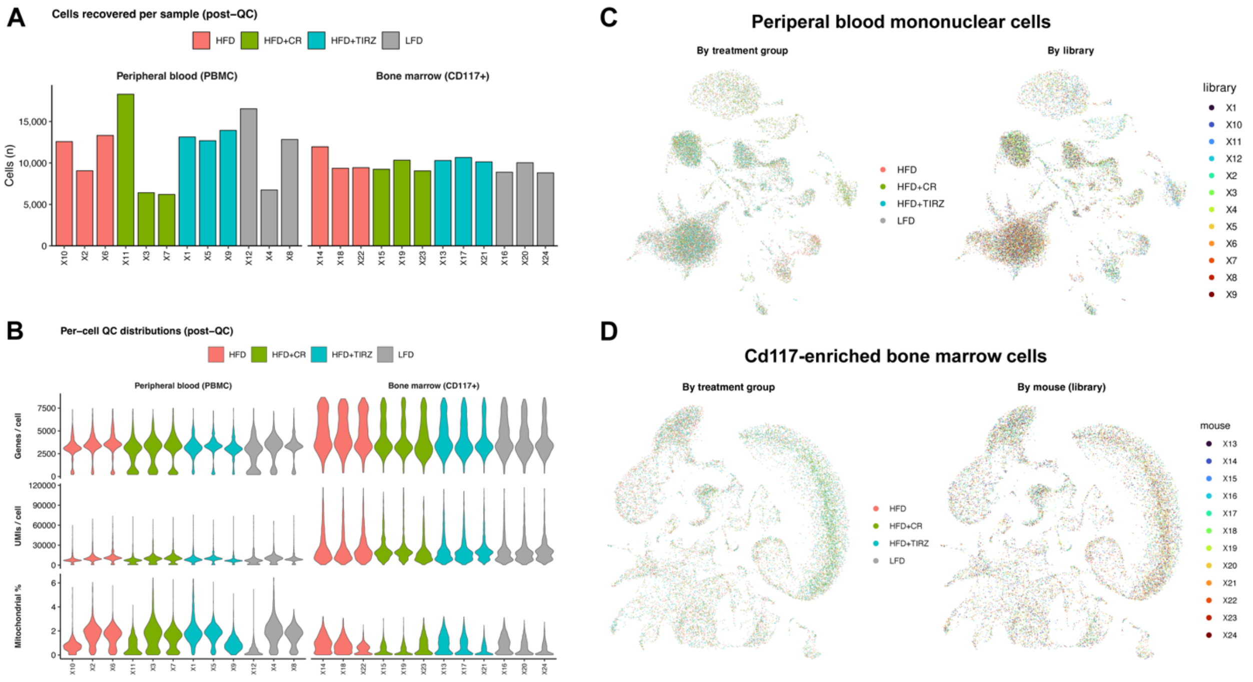


**Supplementary Figure 1: Single-cell sequencing dataset quality control (QC) and integration. (A)** Number of cells recovered per library after QC, for peripheral blood mononuclear cells (PBMCs) and CD117-enriched bone marrow cells (colored by group; 256,555 cells total: 138,622 PBMC and 117,933 CD117$^{+}$ marrow). **(B)** The number of genes per cell, unique molecular identifiers per cell, and the percent of mitochondrial transcripts per cell after QC (colored by group). **(C-D)** Harmony integration by treatment group and by library for **(C)** PBMCs and **(D)** CD117-enriched marrow cells. UMAPs are colored by treatment group (left) and by library (right). Note that PBMC libraries were generated by pooling peripheral blood from 2 mice (n=3 libraries per group; n=6 mice per group). CD117-enriched marrow libraries were generated from one individual mouse (n=3 per group).

**Supplementary Figure 2:**

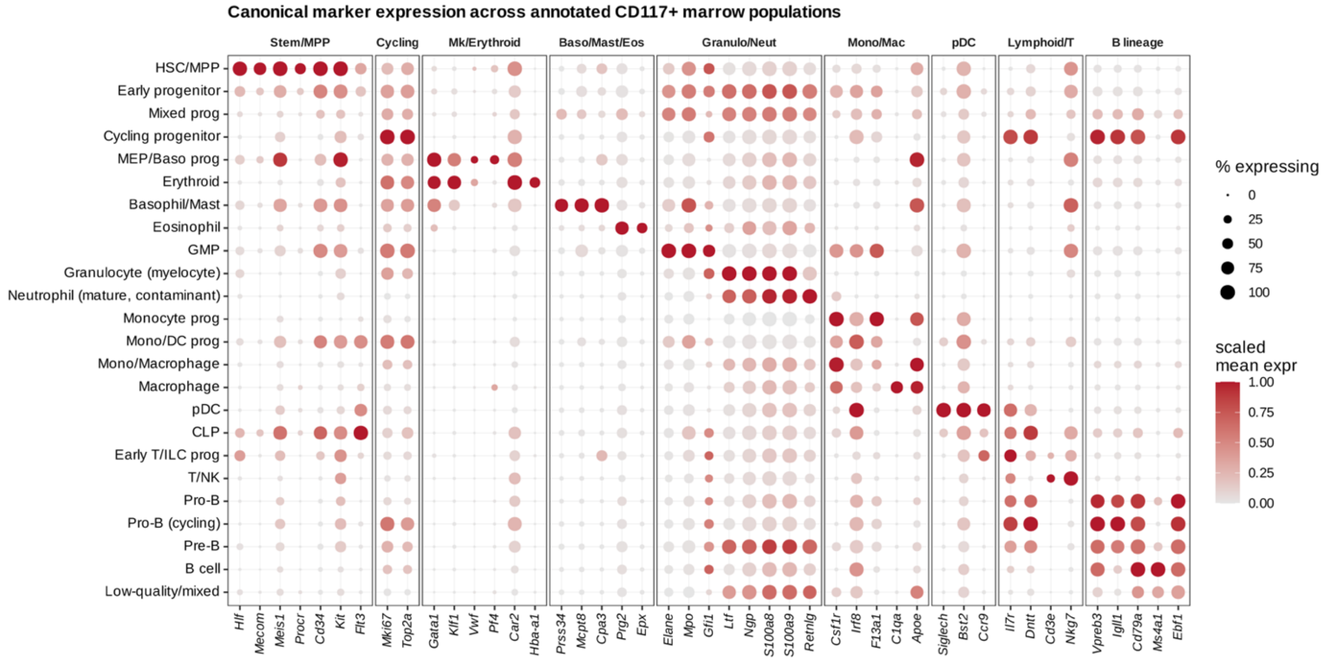


**Supplementary Figure 2: Marker-based annotation of CD117-enriched bone marrow.** Dot plot showing the genes used to define lineages in CD117 (c-Kit)-enriched marrow scRNA-seq experiment (12 libraries; n=3 mice per group). There were 117,933 cells in 41 Leiden clusters (resolution 1.0), which were assigned to 24 hematopoietic populations based on expression of canonical marker genes. Dot size represents the percentage of cells expressing the indicated gene; the color indicates the per-gene scaled expression. Markers were grouped by the lineages they defined.

**Supplementary Figure 3:**

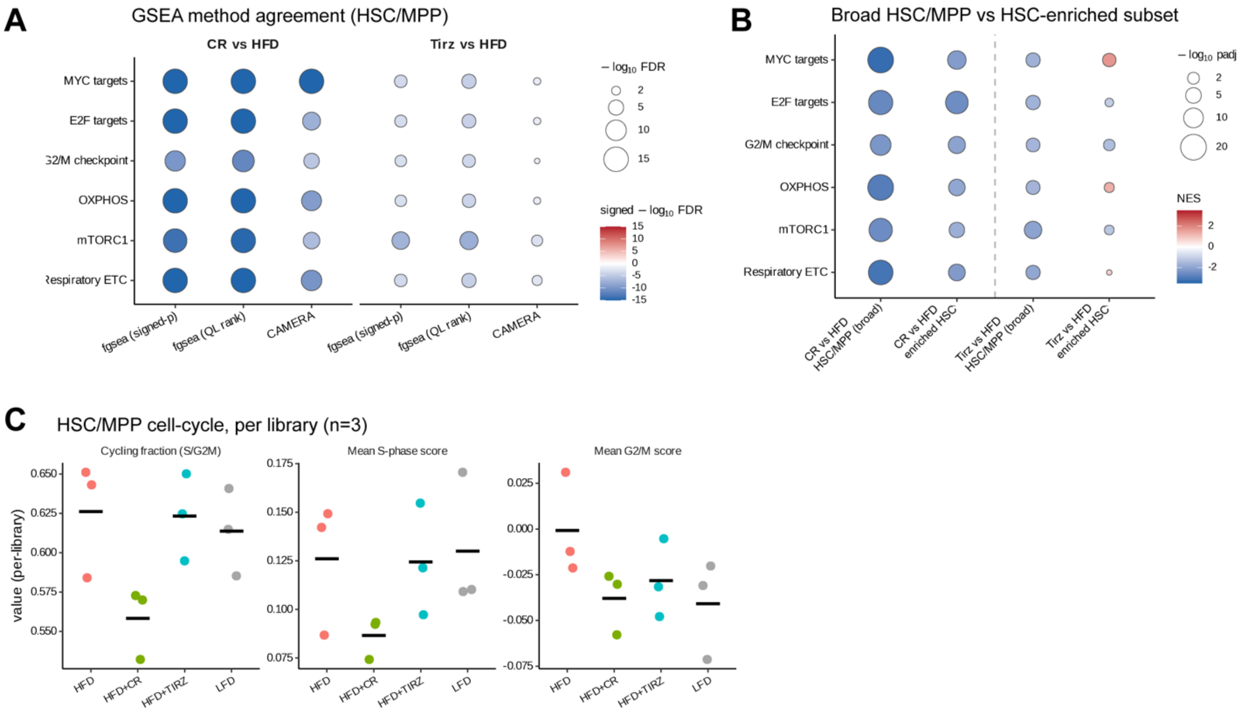


**Supplementary Figure 3: CR-mediated suppression of proliferative and metabolic gene signatures persists in progenitors with alternative enrichment methods. (A)** Indicated pathways for CR vs. HFD, and Tirz vs. HFD scored by three different methods, including 1. fgsea ranked by signed significance, 2. fgsea ranked by the edgeR quasi-likelihood statistic, and 3. CAMERA all showing methodological agreement. Dot size represents the -log10 FDR, while dot color represents the signed −log10 FDR. **(B)** The full HSC/MPP compartment (“broad”) vs. a stringent HSC subset (“core”, top tertile of an HSC-stemness signature) for CR vs. HFD and (left) and Tirz vs. HFD (right). Dot size represents the −log10 adjusted *p*-value, while the color scale is normalized enrichment score (NES). **(C)** HSC/MPP cell cycle by library showing the per mouse cycling fraction scanpy cell-cycle scoring. Symbols are individual libraries/mice (n=3 per group). Bars represent the mean.

### Supplementary Figure 4:

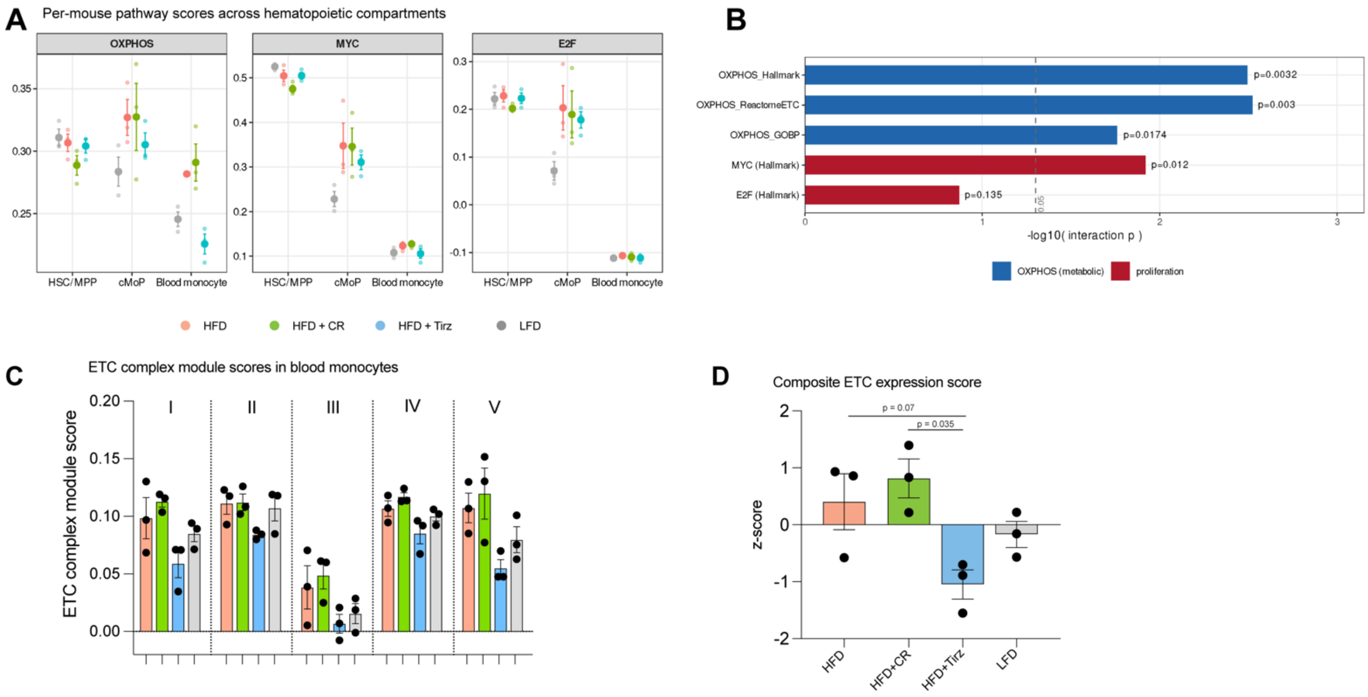


**Supplementary Figure 4: Concordance of tirzepatide-mediated OXPHOS and ETC component suppression across gene set pathway definitions in mature monocytes. (A)** Per-library module scores for OXPHOS, MYC, and E2F across the differentiation spectrum from HSC/MPP, cMoP, to blood monocytes for all four groups (mean ± SEM, while symbols represent individual libraries). **(B)** Treatment-by-compartment interaction for OXPHOS defined using three independent gene sets, together with Hallmark MYC and E2F targets. Bars show −log10(*p*); dashed line indicates *p*=0.05. **(C)** Module scores for individual electron transport chain complexes (complexes I–V) in blood monocytes by indicated treatment group. **(D)** To summarize the coordinated trends across the ETC complexes in **(C)**, a composite ETC expression z-score was calculated, showing reduced composite expression in tirzepatide relative to CR (*p*=0.035). n=3 per group.
